# Colistin resistance genes in *Escherichia coli* isolated from patients with urinary tract infections

**DOI:** 10.1101/2024.01.16.575981

**Authors:** Waleed M. Al Momani, Nour Ata, Ahmed O. Maslat

**Affiliations:** Department of Basic Medical Sciences, Faculty of Medicine, Yarmouk University, 211-63 Irbid, Jordan; Department of Biological Sciences, Faculty of Science, Yarmouk University, 211-63 Irbid, Jordan

**Keywords:** Multi drug resistant, colistin resistance, *Escherichia coli*, Urinary Tract Infections, MCR genes, multiplex PCR, Jordan

## Abstract

**Introduction:** Antimicrobial resistance is alarmingly high because it happens in humans, environment, and animal sectors from a “One Health” viewpoint. Due to the fact, that *Escherichia coli (E. coli)* is broadly disseminated in all sectors, the food web and the environment may have a role in carrying colistin resistance genes from animals to humans. The rise of plasmid-mediated mobile colistin resistance (*MCR*) genes threatens colistin efficacy, which is the last line to remedy Gram-negative infections multidrug resistance (MDR).

**Objectives:** The current study aimed to investigate development of colistin resistance (*MCR* 1-5) genes between *E. coli* isolated from patients with urinary tract infections (UTI) in Jordan.

**Methods:** *E. coli* (n=132) isolated were collected from urine specimens. *E. coli* isolated from human UTI patients were examined for genes resistance to colistin *MCR* (1-5). All isolates were investigated against 20 antimicrobials utilizing the standard disk diffusion method. For analysis of colistin resistance, the broth microdilution technique was employed. In addition, the *MCR* (1-5) genes were detected by multiplex PCR assay.

**Results:** Out of 132 isolates, one isolate was colistin-resistant, having a minimum inhibitory concentration of 8 μg/mL and possessing the *MCR* -1 gene. A total of 132 *E. coli* isolates showed high resistance against penicillin, amoxicillin, cephalexin, nalidixic acid, tetracycline, and cefepime in the percentages of 100%, 79.55%, 75.76%, 62.88%, 58.33%, and 53.79%, respectively. However, resistance was lowest towards fosfomycin (6.06%), florfenicol (10.61%), and chloramphenicol (15.91%). Significant differences were observed between *E. coli* isolated from pediatrics and those isolated from adults.

**Conclusion:** This is the first report on the presence of the plasmid-coded *MCR*-1 gene recovered from *E. coli* from a patient with UTIs in Jordan. That is threatening as colistin is the last line used for infections induced by MDR gram-negative bacteria. There is a crucial need for control and harsh utilization of antibiotics to control and prevent the emergence and prevalence of colistin resistance genes.

**Summary:** *E. coli* isolated from human UTI patients were examined for genes resistance to colistin *MCR* (1-5). This is the first report on the presence of the plasmid-coded *MCR*-1 gene recovered from *E. coli* from a patient with UTIs in Jordan. That is threatening as colistin is the last line used for infections induced by MDR gram-negative bacteria. There is a crucial need for control and harsh utilization of antibiotics to control and prevent the emergence and prevalence of colistin resistance genes. A total of 132 *E. coli* isolates showed high resistance against penicillin, amoxicillin, cephalexin, nalidixic acid, tetracycline, and cefepime in the percentages of 100%, 79.55%, 75.76%, 62.88%, 58.33%, and 53.79%, respectively

## 1. Introduction

Urinary tract infections (UTIs) induced by antibiotic resistant gram negative bacteria (GNB) are the most common bacterial infections faced by clinicians, which is of growing concern due to limited treatment options [1]. Though other bacteria in the *Enterobacteriaceae* family can induce UTIs [2], *E. coli* is the most etiological agent of UTI accounting for up to 80% of all cases [3].

Treatment of UTI is greatly complicated by the emergence of multidrug-resistant (MDR) isolates [4], because of this escalating problem of the prevalent of MDR *E. coli* pathogens, related to the exhausting antibiotic innovation line, colistin has been reused in clinical approaches after being categorized by the World Health Organization (WHO) as one of the antibiotics of crucial significance in human clinical settings [5].

Colistin is regarded as the last-line antibiotic utilized for the therapy of acute infections induced by MDR GNB [6]. Colistin is narrow-spectrum antibiotic that has important action against most partners of the *Enterobacteriaceae* family and common non-fermentative GNB [7].

In 2015, researchers in China reported the presence of the plasmid-mediated colistin resistance *MCR* -1 gene in *E. coli*, which can be transmitted from one bacterium to another and encodes phosphoethanolamine transferase. encodes phosphoethanolamine transferase led to the addition of phosphoethanolamine (a cationic molecule) to lipid A from LPS, which changes the cell membrane charge, and as consequence, colistin (the cation) is unable to attach and induce cell membrane degradation, thus conferring resistance to colistin [8].

There is an idea that *MCR* -1 is derived from animals and then transferred to humans by horizontal transmission. This is because *E. coli* isolates that produce *MCR*-1 have been determined in animal food products [9]. The thoughtless use of colistin in the veterinary sector, particularly in the lack of severe lawmaking has contributed to the global dispersal of the *MCR*-1 gene in 10% of animal isolates and in 0.1–2% of human isolates [10].

Although there are studies that reported the presence of *MCR* genes isolated from patients with the urinary tract in many countries, However, there is no study that investigated the prevalence of *MCR* genes in Jordan from urinary tract patients. In the present study, we aimed to shed light on the happening of colistin resistance among *E. coli* isolated from patients with UTIs in Jordan.

## 2. Methods

### 2.1. Sample Collection and Identification

This study was conducted over the period of 6 months between January to June 2022 and included 132 *E. coli* isolates from the urine cultures of patients with UTI. All participants between the ages of 3 months to 85 years. All isolates were obtained from the Princess Rahma Hospital in Irbid and a clinical diagnostic lab in Amman, Jordan. Samples were streaked onto MacConkey, eosin methylene blue, and blood (Oxoid, UK) agar plates. Following incubation at 37 °C for 24 h, All isolates were confirmed as *E. coli* by standard biochemical tests including IMViC and Kligler Iron Agar tests, and also by Molecular identification by the polymerase chain reaction (PCR) using the Universal Stress Proteins A (UspA) gene with 884 bp band size was used [11]. *E. coli* NCTC 12900 UK was used as the positive control. The study was approved by the ethics committee at Yarmouk university.

### 2.2. Antimicrobial Susceptibility Testing (AST)

The antibiotic susceptibility profile for 132 *E. coli* isolates was determined using the disk diffusion technique on Mueller–Hinton agar (Oxoid, UK) using the suspension equivalent in turbidity to 0.5 McFarland. Then, plates were incubated overnight at 37°C. The results were interpreted according to Clinical Laboratory Standards Institute (CLSI,2017) [12]. The *E. coli* isolates were defined as MDR (resistant to three or more antimicrobial classes) based on the International Expert proposal for Interim Standards Guidelines [13]. The following used antibiotics were tested: Cephalexin (30μg), Penicillin (10 μg), Ciprofloxacin (5 μg), Doxycycline (30 μg), Aztreonam (30 μg), Imipenem (10 μg), Gentamycin (10 μg), Florfenicol (30 μg), Kanamycin (30 μg), Tigecycline (15 μg), Cefepime (30 μg), Amoxicillin-clavulanate (30 μg), Cefoxitin (30 μg), Sulphamethaxazole-trimethoprim (25 μg), Chloramphenicol (30 μg), Tetracycline (30 μg), Fosfomycin (50 μg), Meropenem (10 μg), Amoxicillin (10 μg), Nalidixic-acid (30 μg) (Oxoid, UK).

The minimum inhibitory concentration (MIC) of colistin (colistin sulfate powder, DADvet, Jordan) against the collected 132 *E. coli* isolates was detected by the micro-dilution broth using Muller-Hinton broth (Oxoid, UK). The MICs values ranged from 128 μg /ml to 0.25 μg /ml in a two-fold dilution order. The clinical breakpoints for colistin resistance were defined according to the European Committee on Antimicrobial Susceptibility Testing (EUCAST) and CLSI statements when the MIC value was >2 μg/mL [14]. *E. coli* NCTC 12900 UK was used as the susceptible-control reference strain for disk diffusion and MIC testing.

### 2.3. Detection of the colistin resistance genes by multiplex PCR

DNA was extracted using the boiling method [15], briefly, a 300 μl bacterial suspension was prepared from fresh *E. coli* colonies grown on nutrient agar (Oxoid, UK), the suspension was vortexed and then incubated in a dry bath (Cleaver, UK) at 100°C for 10 minutes followed by immediate incubation on ice for another 10 minutes. After that, samples were placed in a centrifuge (HERMLE, Germany) and centrifuged at full speed for 10 minutes, the supernatant was stored at -20°C and used as a template for PCR.

All *E. coli* isolates (n=132) were screened using multiplex PCR for the presence of mobile colistin resistance genes *MCR* (1-5). The multiplex PCR assay was done according to the European Centre for Disease Prevention and Control [16], the reaction was performed in a volume of 20 μl containing 4 μl of 5x HOT FIREPol® Blend Master Mix (Solis BioDyne, Estonia), 4 μl DNA template,6 μl nuclease-free water,1.2 μl of each primer pair (Table 1).. PCR amplification was done in a Thermocycler (BIO-RAD, USA) with an initial DNA denaturation step at 94 °C, 15 min followed by 25 cycles beginning with 30 s of denaturation at 94°C, 90 s of primer annealing at 58°C, and 1 min of extension at 72 °C. The final extension step was performed at 72 °C for 10 min. Amplified products were visualized by electrophoresis using 2% agarose gel electrophoresis followed by staining with ethidium bromide and were visualized under UV light.

**Table 1.**
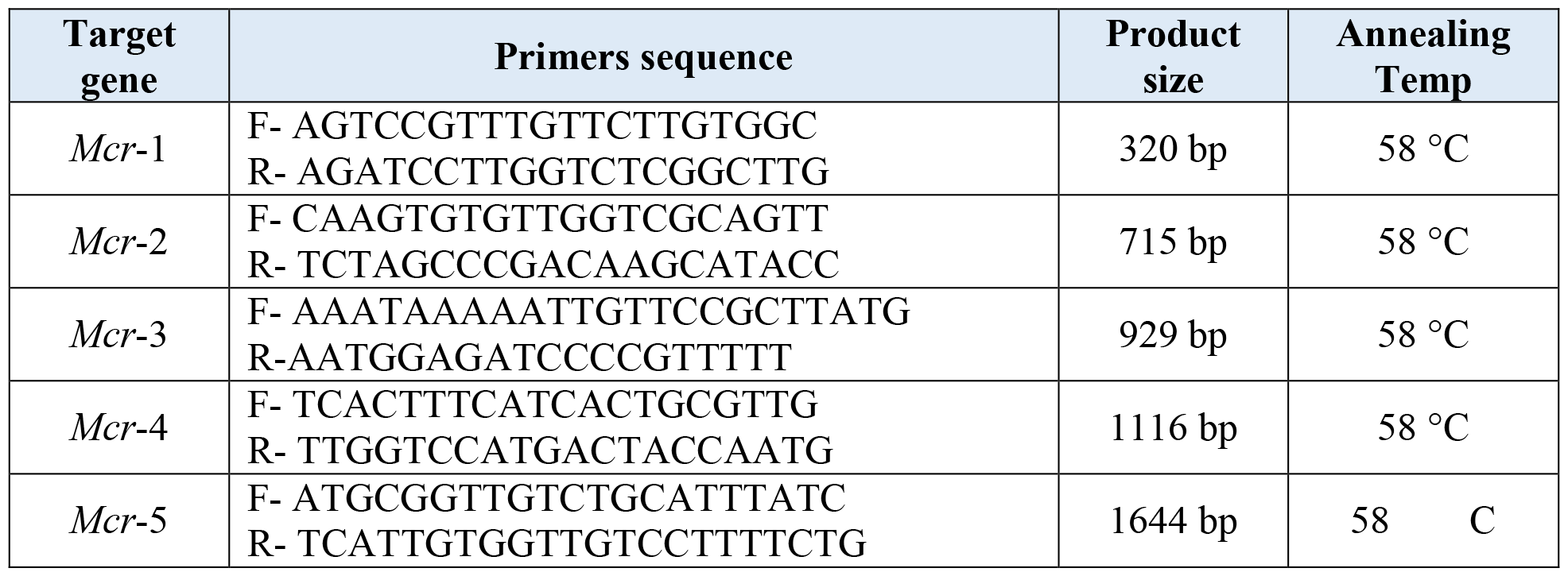
PCR target genes of (*Mcr*1-5), primer sequence, PCR product size, and annealing temperature [17].

## 3. Results

### 3.1. Isolation and characterization of E. coli

Out of 132 urine samples, a total of 132 E. coli were isolated and confirmed, according to gender, 90.2% (n= 119) of the E. coli were isolated from females while 9.8% (n= 13) were isolated from males, according to age 75% (n= 99) of the E. coli were isolated from adults while 25% (n= 33) of the E. coli were isolated from pediatrics. A total of 132 isolates were confirmed as E. coli by PCR amplification, by using Universal Stress Proteins A in Escherichia coli (UspA gene), some of these results are shown in Figure 1

**Figure 1.**
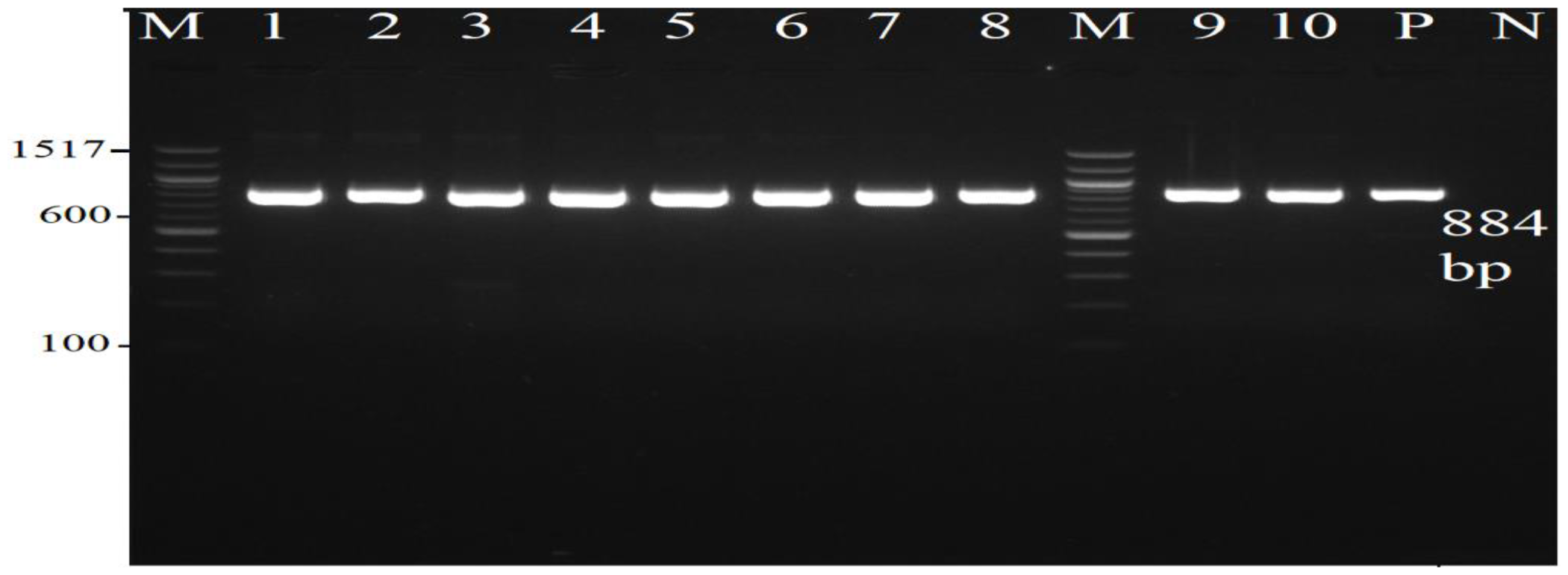
Electrophoresis for *Escherichia coli uspA* gene. Lane M: 100bp ladder; lane 1 to 10: samples; lane P: positive, and N: negative controls.

### 3.2. Antimicrobial Resistance Profiles

A total of 132 isolates showed high resistance against penicillin, amoxicillin, cephalexin, nalidixic acid, tetracycline, and cefepime in the percentages of 100%, 79.55%, 75.76%, 62.88%, 58.33%, and 53.79%, respectively. However, resistance was lowest towards fosfomycin (6.06%), florfenicol (10.61%), and chloramphenicol (15.91%). Reading for each antibiotic was recorded in three categories as resistant (R), intermediate(I), and susceptible(S). The results of the 20 antimicrobials used in this study are shown in (Table 2 and Fig 2).

**Table 2.** Antibiotics susceptibilities for 132 *E. coli* isolates by the disk diffusion method.

| Class of antibiotics |  | Antibiotic tested |  | R | I | S |
| --- | --- | --- | --- | --- | --- | --- |
| 1 | **β-lactams** | 1 | **Penicillin** | 132 (100%) | 0 (0.0%) | 0 (0.0%) |
|  |  | 2 | **Amoxicillin** | 105(79.55%) | 8(6.1%) | 19(14.39%) |
|  |  | 3 | **Aztreonam** | 47(35.61%) | 18(13.64%) | 67(50.76%) |
|  |  | 4 | **Imipenem** | 7(5.30%) | 27(20.45%) | 98(74.24%) |
|  |  | 5 | **Meropenem** | 33(25%) | 17(13%) | 82(62.12%) |
| 2 | **β-lactamase inhibitors** | 6 | **Amoxicillin-clavulanate** | 67(50.76%) | 34(25.76%) | 31(23.48%) |
| 3 | **Tetracyclines** | 7 | **Tetracycline (TE)** | 77(58.33%) | 5(3.79%) | 50(37.88%) |
|  |  | 8 | **Doxycycline (DO)** | 50(37.88%) | 36(27.27%) | 46(34.85%) |
|  |  | 9 | **Tigecycline (TGC)** | 10 (7.58%) | 23(17.42%) | 99(75%) |
| 4 | **Sulfonamides** | 10 | **Sulphamethaxazole-trimethoprim** | 67(50.76%) | 7(5.3%) | 58(43.94%) |
| 5 | **Fluoroquinolones** | 11 | **Nalidixic-acid** | 83(62.88%) | 16(12.12%) | 33(25%) |
|  |  | 12 | **Ciprofloxacin** | 59(44.70%) | 40(30.30%) | 33(25%) |
| 6 | **Aminoglycosides** | 13 | **kanamycin** | 28(21.21%) | 45(34.1%) | 59(44.70%) |
|  |  | 14 | **Gentamycin** | 27(20.45%) | 13(9.85%) | 92(69.70%) |
| 7 | **Cephalosporins** | 15 | **Cefepime** | 71(53.79%) | 15(11.4%) | 46(34.85%) |
|  |  | 16 | **Cefoxitin** | 21(15.91%) | 14(10.61%) | 97(73.48%) |
|  |  | 17 | **Cephalexin** | 100(75.76%) | 0 (0.0%) | 32 (24.24%) |
| 8 | **Phosphoric acid derivatives** | 18 | **Fosfomycin** | 8(6.06%) | 3(2.27%) | 121(91.67%) |
| 9 | **Phenicol** | 19 | **Chloramphenicol** | 21(15.91%) | 9(6.82%) | 102(77.27%) |
|  |  | 20 | **Florfenicol** | 14(10.61%) | 1(0.76%) | 117(88.64%) |

**Figure 2.**
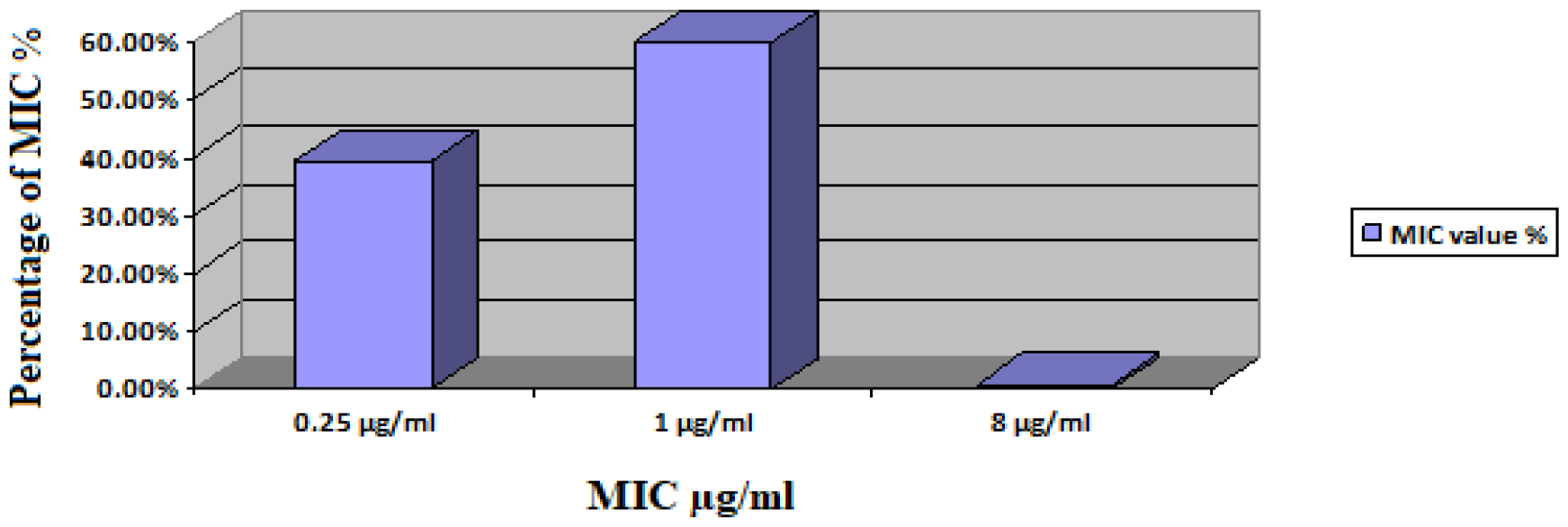
Minimal Inhibitory Concentration value for *E. coli* (n=132) isolates.

To verify the presence of MDR in all *E. coli* isolates, first each isolate was organized according to the number of antibiotics to which *E. coli* exhibited resistance to 20 antibiotics in different classes. Summary of the 132 *E. coli* isolates which resistant to antibiotics shown in (Table 3).

**Table 3.** Summary of the number of *E. coli* isolates (n=132) which resistant to antibiotics (n=20).

| Number of antibiotics that resistance | Number of *E. coli* isolates | Percentage % |
| --- | --- | --- |
| Resistance to 1 antibiotic | 3 | 2.27 % |
| Resistance to 2 antibiotics | 7 | 5.30 % |
| Resistance to 3 antibiotics | 4 | 3.03 % |
| Resistance to 4 antibiotics | 7 | 5.30 % |
| Resistance to 5 antibiotics | 10 | 7.58 % |
| Resistance to 6 antibiotics | 14 | 10.62 % |
| Resistance to 7 antibiotics | 18 | 13.64 % |
| Resistance to 8 antibiotics | 15 | 11.36 % |
| Resistance to 9 antibiotics | 14 | 10.61 % |
| Resistance to 10 antibiotics | 11 | 8.33 % |
| Resistance to 11 antibiotics | 13 | 9.85 % |
| Resistance to 12 antibiotics | 6 | 4.54 % |
| Resistance to 13 antibiotics | 6 | 4.54 % |
| Resistance to 14 antibiotics | 4 | 3.03 % |

Then, each *E. coli* isolate was organized according to the number of classes of antibiotics to which it showed resistance. Summary of 132 *E. coli* isolates which resistant to classes of antibiotics (Table 4). Percentage of *E. coli* isolates that exhibited MDR (88.64%, 117/132).

**Table 4.** Summary of the *E. coli* isolates (n=132) which resistance to classes of antibiotics to show multidrug resistance.

| Number of classes | Number of *E. coli* isolates | Multidrug resistance | Percentage% |
| --- | --- | --- | --- |
| Resistance to 1 class | 5 | No | 3.79% |
| Resistance to 2 classes | 10 | No | 7.57% |
| Resistance to 3 classes | 19 | Yes | 14.4% |
| Resistance to 4 classes | 16 | Yes | 12.12% |
| Resistance to 5 classes | 31 | Yes | 23.5% |
| Resistance to 6 classes | 23 | Yes | 17.42% |
| Resistance to 7 classes | 21 | Yes | 15.9% |

### 3.3. Colistin MIC

Between the 132 strains isolated, a single *E. coli* isolates harbored the *MCR*-1 gene and showed resistance to colistin (MIC =8 μg /ml), The remaining isolates were sensitive to colistin with MIC values < 2 μg/ml. (Table 5, Figure 1).

**Table 5.** Minimal Inhibitory Concentration (MIC) value for132 *E. coli* isolates.

| MIC value | MIC =0.25 µg/ml | MIC =1 µg/ml | MIC =8 µg/ml |
| --- | --- | --- | --- |
| **Susceptibility** | 52(39.39%) | 79(59.85%) | - |
| **Resistance** | - | - | 1(0.76) |

### 3.4. Molecular Identification of colistin Resistance Genes

A total of 132 isolates were screened for the presence for (*MCR* 1, *MCR* 2, *MCR* 3, *MCR* 4, and *MCR* 5) by multiplex PCR. Our results show that 1 of 132 *E. coli* isolates carry *MCR*-1(Fig 3).

**Figure 3.**
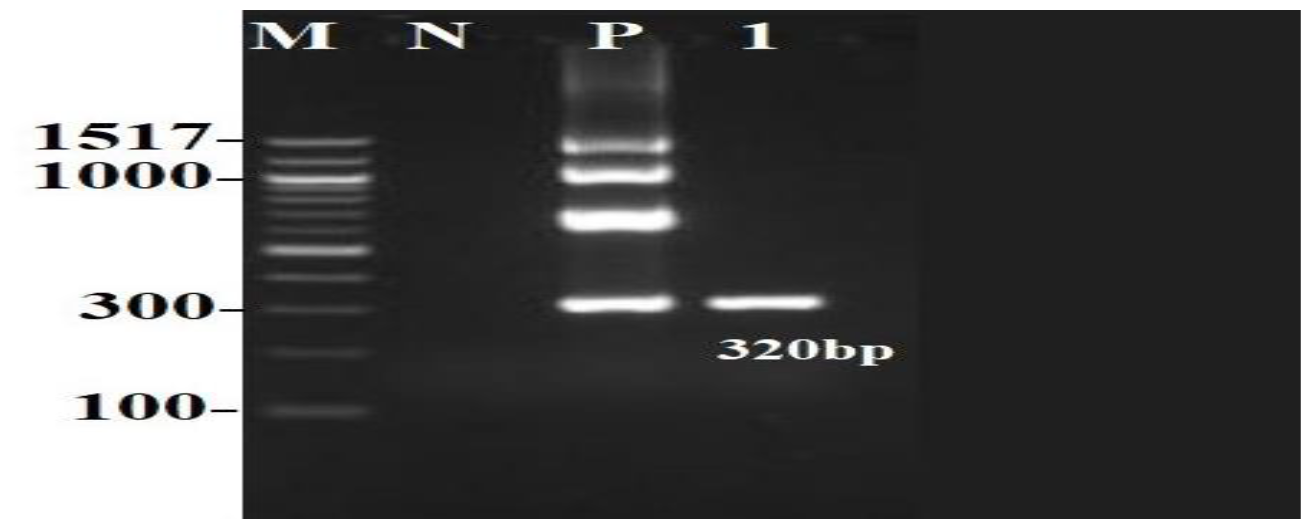
Electrophoresis for a single *E. coli* isolate carries *MCR*-1gene.

### 3.5. A single E. coli isolates harbored the MCR-1 gene

A single *E. coli* isolate harbored the *MCR-*1 gene was deemed to be resistant to colistin with minimum inhibitory concentrations MICs, and multiplex PCR. which exhibited MICs value equal to 8 μg/ml. Summary results for single *E. coli* isolate that show resistance to colistin (Table 6).

**Table 6.** Summary results for single *E. coli* isolate that shows resistance to colistin.

| Source | Age | No. of isolates | *MCR* gene | MIC of colistin (µg/ ml) | Resistant antibiotics | Number of classes |
| --- | --- | --- | --- | --- | --- | --- |
| Human UTI | 27 | 1 | *MCR* -1 | 8 | AMC, P, CIP, ATM, FEP, CL, SXT, C, TE, NA | 7 |

## 4. Discussion

*E. coli* was isolated from urine samples (n= 132), out of which 90.2% (n= 119) originated from females with UTI while 9.8% (n= 13) originated from males, moreover and according to age 25% (n= 33) originated from pediatrics while 75% (n= 99) originated from adults. Urinary tract infections (UTI) display one of the numerous major sex differences between infectious diseases [18]. The structural difference in the female urinary system donates to the development of UTIs [19]. An assumption to clarify this difference is that anatomical disparity, like the short space between the anus and the urethral opening in females or a long urethra in males [18].

*E. coli* is the most etiological agent of UTI accounting for up to 80% of all cases [3, 20]. In this study, *MCR*-1 was detected in one isolate (out of 132, 0.76 %) derived from a urine sample of a 27-year-old female inpatient with a UTI. This prevalence rate was approximately similar to rates reported in Myanmar (0.23%) [21], and Switzerland (0.12%) [22]. Despite its low spread, the existence of *MCR*-1 in a clinical isolate in Jordan could be considered as a sign of start of transmission of resistance genes mainly *MCR*-1 in *E. coli* in humans. On the other hand, higher prevalence rates occurred in Egyptian studies with a prevalence rate of 22*%*, [23] 5.6 *%* [24] and 4.5 *%* [25], and another study on UTIs from Bangladesh with a prevalence rate of 3.52% [26]. Large and improper use of antibiotics creates selective pressure, pursued by the fast rise and outbreak of MDR *Enterobacteriaceae* [24].

This study underlines the prevalence of MDR in urinary *E. coli*. 117 of 132 isolates showed resistance to at least three classes of antibiotics to be multidrug-resistant, and the percentage of *E. coli* isolates that exhibited MDR (88.64%, 117/132). This finding is supported by other similar UTI studies that showed high resistance rates to the commonly used antibiotics [23]. As a potential justification for the presence of such MDR high rate of MDR in UTI patients because *E. coli* inducing UTIs are known for their capability to form biofilms that induce recurrent leading to continuous and resistant infection [27].

Colistin has been more used internationally as an antibiotic of resort for infections resulting from Gram-negative bacteria [9]. In addition, since the first report in China in 2015, the incidence of plasmid-mediated colistin resistance gene *MCR*-1 has been identified in *Enterobacteriaceae* from animals and humans in different countries [28]. The fluctuation in colistin resistance among different studies can be explained by the number of cases, the general situation of patients, geographical regions, different antibiotic regulations, and compliance with infection management measures [24]. The mobile colistin resistance *MCR*-1 gene is more common than other *MCR* genes, a result that is supported by our study as well as previous researches [21-25]. On the other hand, *MCR*-2 gene was reported in *E. coli* isolated from patient with UTI in a study conducted in Bangladesh [26].

All *E. coli* isolates were resistant to penicillin. In addition, highest resistance rates, surpassing 50%, were detected for amoxicillin-clavulanate, cephalexin, cefepime, tetracycline, amoxicillin, nalidixic-acid, and sulphamethoxazole-trimethoprim. Moreover, high susceptibility rates, exceeding 75%, were detected for florfenicol, tigecycline, chloramphenicol, and fosfomycin. these resistance rates are approximately similar to rates studied in Egypt demonstrating that *E. coli* isolates from patients with UTIs were highly resistant to amoxicillin-clavulanate, nalidixic-acid, sulphamethoxazole-trimethoprim, tetracycline and cefepime [25].

In this research, the highest resistance rates have been found against β-lactams, a possible reason regarding the excessive resistance rate to those antimicrobial agents that is in Jordan, and within the previous few years (2012–2015) an excessive rate of ESBL-producing *E. coli* (43–54%) has been isolated from UTIs patients with which is drastically higher than what was reported in 2009 (10.8%) [29]. Highest resistance to these antibiotics may be also because of doctors’ empiric antimicrobial prescription, self-prescribing, non-obligation, and drug consumption without permission of the doctor [30].

The detection of one isolate *MCR*-1 gene in *E. coli* urinary tract strains was proved by Multiplex PCR similarly, *MCR*-1 was shown in colistin-resistant *E. coli* isolates from China [31]. Single *E. coli* isolate that resisted colistin, was also resisted multiple classes of antimicrobials, but was susceptible to gentamicin, florfenicol, kanamycin, tigecycline, and fosfomycin. Multiple studies have shown that colistin-resistant isolates exhibit high resistance to multiple classes of antimicrobials [24].

In the current study, the isolate was deemed resistant to colistin also with minimum inhibitory concentration. This *E. coli* isolate carrying the *MCR* -1 gene exhibited MIC value equal to 8 μg/ml (MIC > 2 μg /ml). These results are similar to studies in Egypt, and Saudi Arabia [25]. Among the 132 strains isolated, a single *E. coli* isolates harbored the *MCR-*1 gene and showed resistance to colistin (MIC =8 μg /ml), The remaining isolates were sensitive to colistin with MIC values < 2 μg/ml. These results are similar to studies in Egypt [23, 25]. On the other hand, other studies showed that some negative isolates in PCR exhibited phenotypic colistin resistance in MIC [24, 26].

The MIC values for resistant isolates ranged from 2-128μg/ml, and 23 (6.4%) isolates exhibited MIC values of ≥ 8 [32]. In addition, *E. coli* isolates from broiler displayed the highest resistance against tetracycline 360 (100%), penicillin 359 (99.7%), and amoxicillin 357 (99.2%) [32]. The worldwide increased spread of *MCR*-1 among animal isolates in comparison with human clinical isolates implies that animals are probable sources of *MCR*-1 in humans. Furthermore, the misusage of colistin in agriculture and the poultry sector may be the principal reason for the elevated prevalence of *MCR*-1 in bacteria isolated from animals and animal yields [33]. In veterinary remedies, colistin is excessively used for various purposes, consisting of prophylaxis and treatment of enteric infections as well as administration with meals in poultry farms to save infections caused by pathogenic bacteria [23].

This study provided data about the antimicrobial resistance pattern in *E. coli* isolated from patients with UTIs. The study concentrated on UTIs because they still constitute the major source of infection for humans beings [22], *E. coli* from community-received infections are at the interplay among the environment and hospitals, playing a possible part as an exchange for *MCR*-like genes within the environment [22], and *E. coli* is the most common member of *Enterobacteriaceae* isolated from the clinical samples [24].

The current study had several fundamental limitations such as the study was performed via a cross-sectional design without future follow-up because of resource limitations, and recurrent infections were not sequestered from first-time infections. The data in the current study exhibit a worrisome spread of the colistin-resistant *E. coli* carrying *MCR*-1, as those were found in humans and broilers in previous studies in Jordan [32].

The results may reflect the prevalence of colistin-resistant *E. coli* through Jordan, or the silent spread of this gene might happen in humans. Furthermore, analysis of the genetic data of the *MCR*-1-positive strains could help us to comprehend the origin of this gene. The *MCR* gene’s presence in this study infers a massive public health danger for colistin antibiotic as a last-line drug. The *MCR* genes are plasmid-mediated, that can spread by horizontal gene transfer to other commensal and pathogenic bacteria [34, 35]. A coordinated strategy for the deterrence of *MCR*-1 dissemination is needed to restrict the spread of those multidrug-resistant isolates between patients [24]. More forceful rules must be enforced to stop the further dissemination of these colistin resistance genes [26].

## 5. Conclusions

Colistin is taken into consideration as one of the last lines of therapy that had been used to deal with extreme infections resulting from MDR pathogens. The development of plasmid-mediated colistin resistance in *E. coli* is presently an essential problem due to the increased possibility of their prevalence in medical settings. The *MCR*-1 gene prevalence on most continents has been noticed in numerous bacterial isolates from animals, human beings, and the environment, involving *E. coli*. In Jordan, colistin-resistant *E. coli* harboring *MCR*-1 was recorded here in this study for the first time in patients with UTIs. This is concerning and highlights the potential health risks that plasmid-mediated colistin-resistant genes in *E. coli* can pose to millions of humans in Jordan. In addition, guidelines should be carried out on the usage of colistin in human and animal sectors to ensure the success of the therapy and to prevent spread of these resistance genes.

